# Linking Enzyme Upregulation to Autophagic Failure: A Potential Biomarker for GM1 Gangliosidosis

**DOI:** 10.1101/2020.10.28.359083

**Authors:** Sarah Smith, Jessica Larsen

## Abstract

With an increasing aging population, neurodegenerative diseases are having an increased impact on society. Typically, these diseases are diagnosed significantly past symptom onset, decreasing the possibility of effective treatment. A non-invasive biomarker and specific target are needed to diagnose and treat the disease before late-stage symptoms. GM1 Gangliosidosis is a lysosomal storage disease where lysosomal enzyme β-galactosidase is missing. As a result, GM1 ganglioside is not broken down and accumulates in the cell, ultimately leading to cell death. One of the main aspects of GM1 Gangliosidosis, and other neurodegenerative diseases, is impaired autophagy: reduced fusion of autophagosomes and lysosomes to degrade cellular waste.

In this paper, we show that healthy cells (NSV3) have approximately 13 times more co-localization of lysosomes and autophagosomes than GM1 Gangliosidosis-diseased cells (GM1SV3), as demonstrated via immunofluorescence. GM1SV3 fold normal enzyme activity of β-galactosidase was downregulated while mannosidase, and hexosaminidase A were both upregulated. When inducing impaired autophagy in NSV3 via starvation, co-localization gradually decreases with increased starvation time. Most notably, after 48-hour starvation, healthy cells (NSV3) showed no significant difference in co-localization compared to GM1SV3. NSV3 under starvation conditions showed a significant increase between time starved and fold normal enzyme activity, with a positive correlation being observed. Activities of mannosidase, and hexosaminidase A of starved NSV3 closely resemble, and surpass, GM1SV3 after 12-hour starvation.

These observations have the potential to expand the conversation regarding impaired autophagy as a potential biomarker for disease progression and diagnostics and as a treatment target.

## 1. Introduction

As the aging population is increasing, neurodegenerative diseases are having an increasing impact on society, growing from a ~6% to a ~11% global burden in less than five years^1^. The development of treatments for neurodegeneration is currently suffering from lack of early diagnosis. Because diagnosis typically occurs well past symptomatic presentation, treatments are ultimately meant to relieve some of the more detrimental symptoms but have limited capabilities of repairing already damaged cells. Diseases of this kind include, but are not limited to, Alzheimer’s, Parkinson’s, Huntington’s, and lysosomal storage diseases (LSDs). Many of these disorders are beginning to be thought of and categorized together as diseases of the lysosome^2–7^. Literature in multiple neurodegenerative diseases show that lysosomal hydrolases are upregulated early in neurodegeneration, but the relationship between enzyme upregulation and neurodegenerative pathways remains unknown^8,9^. In the last decade, researchers have been looking for lysosomal-oriented biomarkers to explain this behavior and diagnose disease, including changing activities of β-galactosidase and β-hexosaminidase in Alzheimer’s related dementia^10^ and reduced β-glucocerebrosidase coupled with increased β-hexosaminidase activity Parkinson’s disease^11^. The general pathological changes that occur in the brain during neurodegeneration are widely known and accepted. Early stages of disease are marked by accumulation of autophagosomes, leading to ultimate cell death through known pathways (Figure 1). Despite this, the links between pathology and disease presentation on an individual disease basis remains unknown^8,12–15^. As the field begins to search for the link between disease progression and lysosomal function, non-invasive disease biomarkers may be identified.

**Figure 1.**
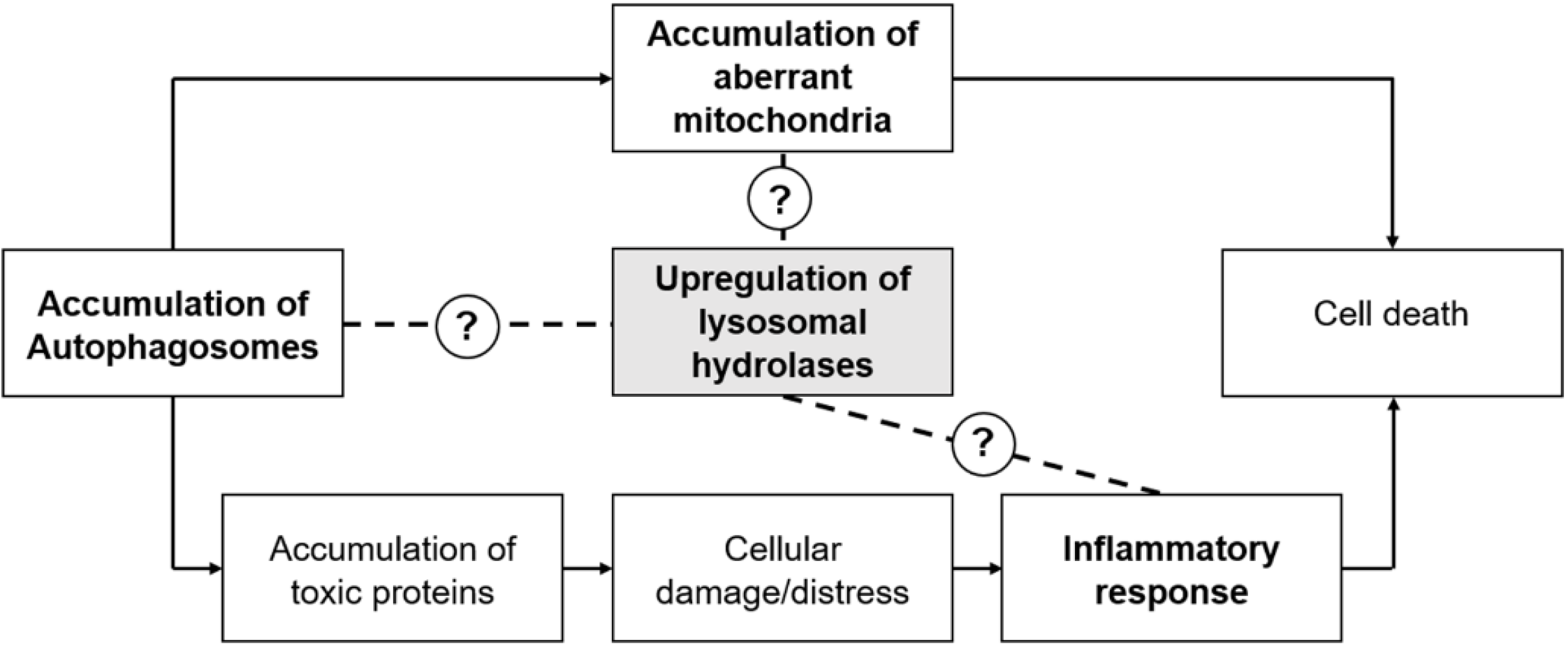
Pathway of neurodegeneration, beginning with accumulation and autophagosomes and leading to ultimate cell death. The role of upregulated lysosomal hydrolases remains unknown, as indicated in the figure. This paper explores a potential link between accumulation of autophagosomes and hydrolase upregulation.

Finding non-invasive biomarkers for brain diseases that can aid in earlier diagnosis can be extremely challenging due to the isolation of the brain from the body through the blood-brain barrier (BBB). However, there are a few non-invasive biomarkers used in diagnostics today. Traditionally cerebrospinal fluid has been used to diagnose Alzheimer’s Disease by measuring levels of total tau (T-tau), tau phosphorylated at threonine 181 (P-tau_181P), and amyloid-B 1-42 (AB_1-42) with other options including miRNAs and neuroinflammatory markers^16,17^. Yet, these methods are not fully reliable due to sample degradation and natural marker levels changing with age. Parkinson’s Disease has no definitive diagnostic test, with medical professionals having to rely on symptom presentation, brain scans and ultrasounds^18,19^.

This paper focuses on the connection between disease progression and lysosomal enzyme activities in an LSD, GM1 Gangliosidosis. In GM1 Gangliosidosis, the lysosomal enzyme β-galactosidase (βgal) is missing or underproduced^7,20^. βgal has the primary role of catabolizing down GM1 ganglioside to GM2 ganglioside^21^. In the absence of this enzyme, GM1 ganglioside is not broken down and accumulates in the cell with other storage products, which leads to swollen lysosomes and ultimate cell death. The lack of βgal leads to upregulation of other lysosomal enzymes, mannosidase and hexosaminidase A^22^. This is believed to be a compensatory mechanism, aiding the cell in digestion of byproducts normally broken down by βgal. Although it is accepted that this upregulation occurs, the point within the disease progression where lysosomal enzyme upregulation occurs is unknown. Further, understanding this relationship may aid in the identification of enzyme activity levels that could be used as non-invasive biomarkers of disease throughout progression.

Neurodegeneration usually occurs in three stages: impaired autophagy, accumulation of dysfunctional mitochondria, and neuroinflammation (**Error! Reference source not found.**). This paper focuses on elucidating the relationship between autophagy and lysosomal hydrolase upregulation. In healthy cells, the autophagic process breaks down unwanted cell-matter. An autophagosome envelops the unwanted material and fuses with a lysosome. The lysosomal enzymes then breakdown the material to be recycled by the cell to make new molecules. However, one way that this process is impaired, which is believed to occur in GM1 gangliosidosis^23^, is failing autophagosomal-lysosomal fusion, causing a buildup of both organelles (Figure 2). This, in turn, leads to the accumulation of toxic proteins or swollen lysosomes in the autophagosomes that cannot be broken down. Here, we present our analysis of the relationship between autophagic failure and lysosomal hydrolase upregulation in a cellular model of GM1 Gangliosidosis, isolated fibroblasts from GM1-gangliosidosis affected felines.

**Figure 2.**
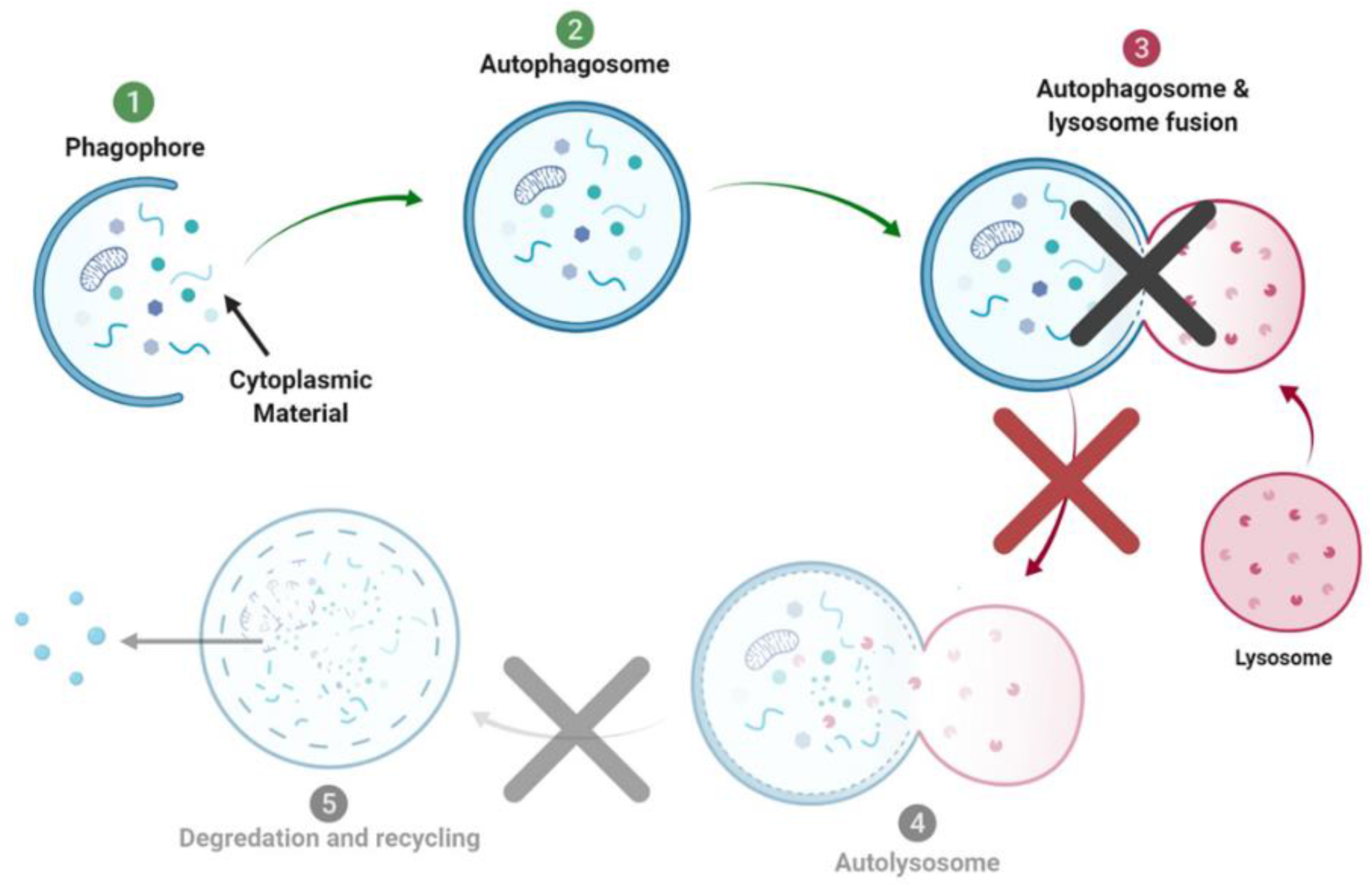
Autophagic Process during Neurodegeneration. In many neurodegenerative diseases, lack of autophagosomal-lysosomal fusion is believed to be the main cause of autophagic failure and apoptosis.

## 2 Materials and Methods

### 2.1. Materials

Dr. Douglas Martin of Auburn University donated skin fibroblasts from healthy cats, NSV3 cells, and GM1 gangliosidosis-affected cats, GM1SV3 cells. Media was Dulbecco’s Modified Eagle Media (DMEM) (ThermoFisher, Cat. Num. 12430, Waltham, MA) with 10% Fetal Bovine Serum (FBS) and 1X Penicillin-Streptomycin (PS) (ThermoFisher, Cat. Num. 15140122, Waltham, MA). Passages were performed using 0.25% Trypsin (Corning, Cat. Num. 25-053-Cl, Manassas, VA). CellLight™ Lysosomes-GFP, BacMam 2.0 (ThermoFisher, Cat. Num. C10507, Waltham, MA) and LC3 staining with an anti-LC3B Rabbit Polyclonal Antibody (ThermoFisher, Cat. Num. PA1-46286, Waltham, MA) (1:300) and Donkey anti-Rabbit IgG (H+L) ReadyProbes™ Secondary Antibody conjugated to Alexa Fluor 594 (ThermoFisher, Cat. Num R37119, Waltham, MA were used for immunofluorescence. Fixation occur with 4% paraformaldehyde (PFA) (Ted Pella, Cat. Num. 18505, Redding, CA) or glutaraldhyde (Sigma Aldrich, Cat. No. G7651-10ML, St. Louis, MO) and growth/starvation media. Cells were permeabilized with 0.1% Triton 100X (Acros Organics, Cat. No. 215682500, Waltham, MA). When blocking was required, 5% donkey serum (Jackson ImmunoResearch, Cat. No. 017-000-121, West Grove, PA) was used. Mounting medium, Vectashield Antifade Mounting Medium with DAPI (Vector, Cat. Num. H-1200, Burlingame, CA) and cytoseal xyl (Fisher Scientific, Cat. No. 22050262, Waltham, PA), were also used.

Enzyme assays and x-gal staining require the use of the following materials: 4-methylumbelliferone (4-MU) (Sigma Aldrich, Cat. No. M1381, St. Louis, MO), 10N NaOH (BDH, Cat. No. BDH7247-1, Radnor, PA), citric acid (Fisher Scientific, Cat. No. A104-3, Hampton, NH), sodium phosphate (Sigma Aldrich, 7782-85-6, St. Louis, MO), sodium chloride (BDH, Cat. No. BDH9286, Radnor, PA), 0.1% Triton X-100 (Acros Organics, Cat. No. 215682500, Waltham, MA), magnesium chloride hexahydrate (Sigma Aldrich, Cat. No. M2670-100G, St. Louis, MO), sodium deoxycholate (Sigma Aldrich, Cat. No. D6750-10G, St. Louis, MO), potassium ferricyanide (Sigma Aldrich, Cat. No. 702587-50G, St. Louis, MO), potassium ferrocyanide (Fisher Scientific, Cat. No. P236500, Waltham, PA), and 0.05% bovine serum albumin (Fisher Scientific, Cat. No. BP1600-100, Hampton, NH).

Substrates to measure enzyme activity were 4-methylumbelliferyl β-gal (Acros Organics, Cat. No. 337210010, Waltham, MA), 4-methylumbelliferyl a-D-manopyranoside (Toronto Research Chemicals, Cat. No. M334745, Ontario Canada), 4-methylumbelliferyl 6-sulfa-2-acetoamido-2-deoxy-b-D-glucopyranosidase (Toronto Research Chemicals, Cat. No. M335000, Ontario Canada), and 4-methylumbelliferyl N-acetyl b-D-glucosaminide (Sigma Aldrich, Cat. No. M2133, St. Louis, MO).

Lowry solution A consisted of copper sulfate (Fisher Scientific, Cat. No. C493-500, Waltham, MA), sodium potassium tartrate (BDH, Cat. No. BDH9272, Radnor, PA), and sodium carbonate (BDH, Cat. No. BDH9284, Radnor, PA). Solution B was made of Folin and Ciocalteu’s phenol reagent (Sigma Aldrich, Cat. No. F9252, St. Louis, MO) and MILLI-Q water.

### 2.2. Methods

#### 2.2.1. Healthy and Starvation-inducing Cell Culture

NSV3 cells and GM1SV3 cells were grown in T75 cell-culture treated flasks at 37 °C, 5% CO_2_. Cells were passaged using standard protocols. Starvation media comprised of DMEM, with no Fetal Bovine Serum, and 1X PS (99 units/mL Penicillin, 99 ug/mL Streptomycin) was used to induce autophagy.

#### 2.2.2. Immunofluorescence Analysis of Autophagosomal Behavior

NSV3 and GM1SV3 cells were passaged into chamber slides at a seeding density of 9.35×10^4^ cells per well in normal growth media with 2 uL of CellLight per well. After overnight incubation, cells were fixed with ice cold 4% PFA in the appropriate growth media (normal growth media or starvation media) for 15 minutes. Cells were then permeabilized with 0.1% Triton X-100 in PBS, 3 times for 5 minutes each. A blocking step was performed with 5% donkey serum in PBS for 1 hour. Cells were then washed with PBS and incubated overnight at 4° Celsius with 1:300 primary antibody dilution in PBS supplemented with 4% FBS, for a total working volume of 500 uL. After a second overnight incubation, cells were washed 4 times with PBS for 10 minutes at a time. Secondary antibody was added to the appropriate wells ensuring untreated, primary-only, and secondary-only wells were run during each experiment as internal controls with 500 uL of appropriate growth media and allowed to incubate in the dark at room temperature for 30 minutes. Cells were washed with PBS, chambers removed and 1 drop Vectashield Antifade Mounting Medium with DAPI was added. Slides were then sealed and allowed to dry overnight at 4°Celsius. Images were analyzed using ImageJ and the “Color Inspector 3D” plugin. In order to quantify yellow and blue pixels, the LAB color code was used to isolate the color threshold. Scale bars were specified in the color range for quantification; values used are found in Table 2. After setting the color range, each cell body was isolated and the frequency of yellow and blue pixels was determined. After obtaining this data, the percent co-localization was calculated as the number of yellow pixels per number of blue pixels.

**Table 1.**
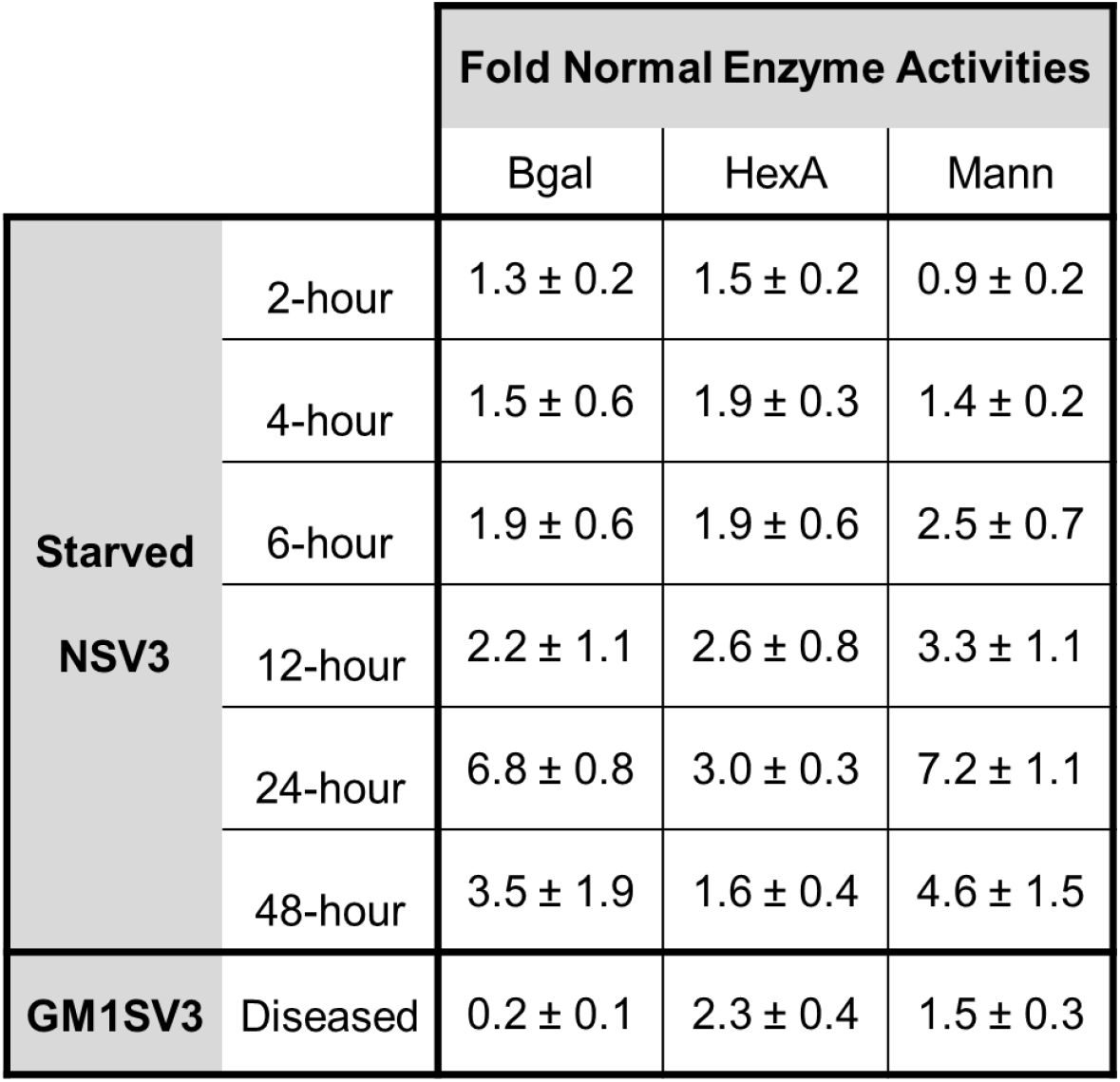
Fold Normal Enzyme Activity Values. Fold normal enzyme activities are measured by comparing activity levels to NSV3 (healthy) cells. Fold normal activity levels in starved NSV3 cells are compared to diseased GM1SV3 cells.

**Table 2.**
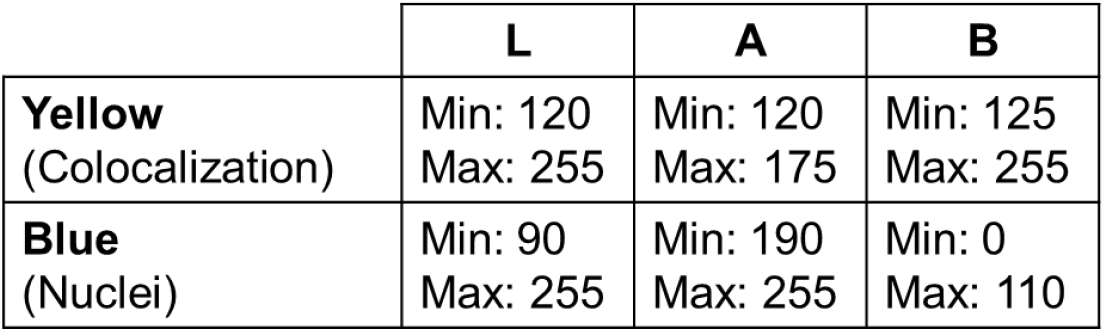
Values used to quantify co-localization in ImageJ

#### 2.2.3. Enzyme Assays

After incubation of NSV3 and GM1SV3 cells in 6-well plates, the media was changed and wells were manually scraped. After centrifugation, cells were re-suspended in Enzyme Isolation Buffer (0.1% Triton X-100 in 50 mM citrate phosphate buffer, pH 4.4 and 0.05% bovine serum albumin) and disrupted using 3 mL syringes. After enzyme isolation, enzyme activity of lysosomal hydrolases β-galactosidase, hexosaminidase A, hexosaminidase B, and mannosidase were measured using 4-methylumbelliferone enzyme assays described previously^49^

#### 2.2.4. X-gal Staining

NSV3 and GM1SV3 cells were passaged into chamber slides at a seeding density of 9.35×10^4^ cells per well in normal growth media. After overnight incubation, x-gal staining was completed as previously described and compared to clinical feline data already published^50^.

#### 2.2.5. Statistical Methods

Statistical analysis between NSV3 co-localization and GM1SV3/NSV3 under starvation conditions was completed by using a two-tailed t-test with samples of unequal variance (p < 0.05). Statistical analysis between NSV3 and GM1SV3 enzyme activity, between NSV3 and NSV3 under starvation conditions enzyme activity, and GM1SV3 and NSV3 under starvation conditions was completed by using a two-tailed t-test with samples of unequal variance (p < 0.05 & p < 0.025).

## 3. Results

### 3.1. Phenotypic Behavior

Phenotypically, NSV3 (healthy) and GM1V3 (affected) cell lines appear the same, with extended, spiny processes typical of fibroblast cell lines, as seen under bright field (BF) (Figure 3).

**Figure 3.**
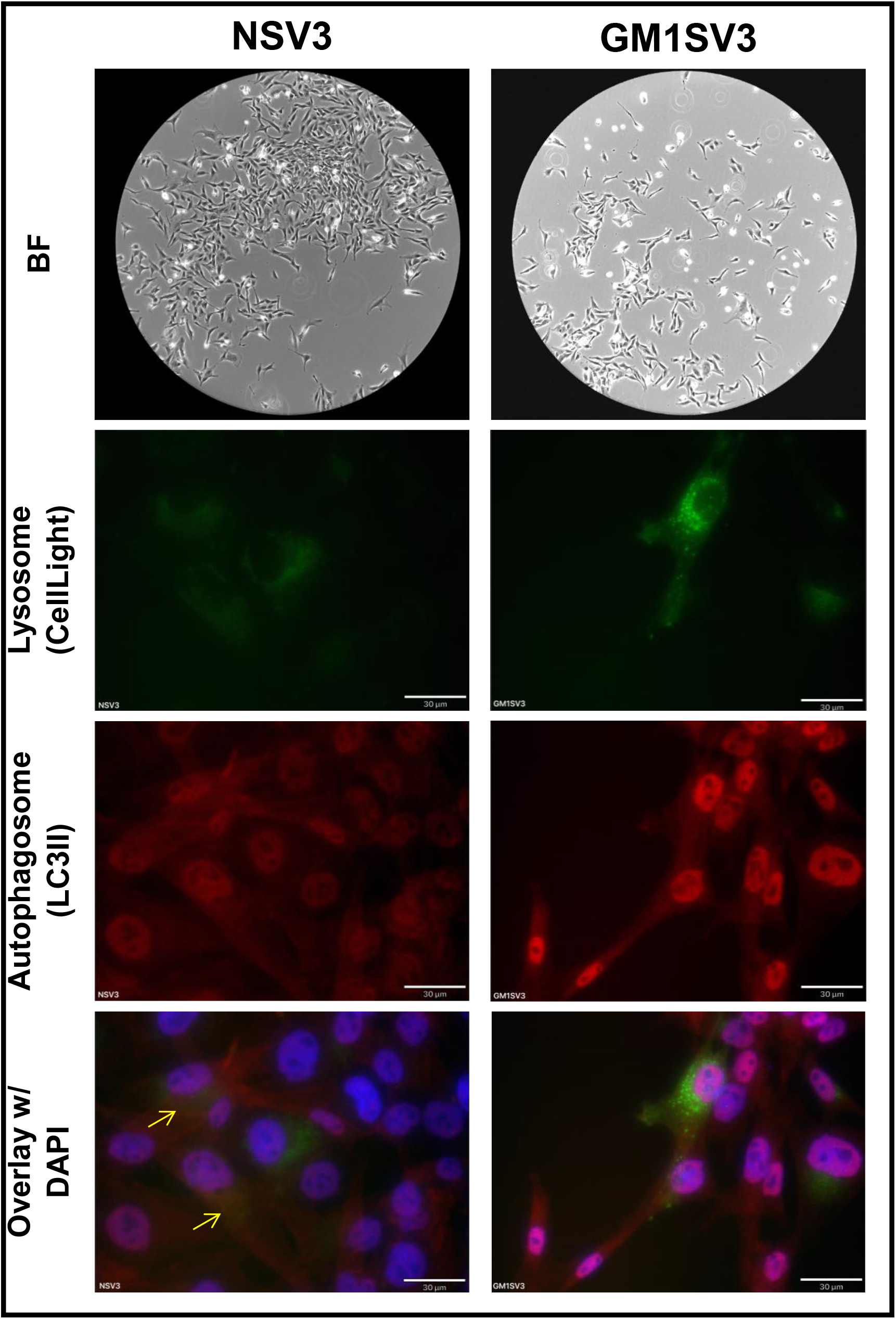
Phenotypic, Lysosomal, and Autophagic behavior of unaltered NSV3 and GM1SV3 cell lines.

### 3.2. Autophagic Behavior

Differences are seen after transfection with CellLight GFP-Lysosome and immunofluorescence with Anti_LC3B. The intensity of the green fluorescence, indicative of lysosomes, in GM1SV3 cells was much greater than NSV3 samples. Similar behavior was observed when looking at red fluorescence with increased red intensity, indicative of autophagosomes, in GM1SV3 when compared to NSV3. When overlaying the green and red fluorescent channels, yellow coloring, indicative of co-localization of green and red fluorescence, is present in NSV3 samples and negligible in GM1SV3 samples (Figure 3).

### 3.3. Characterization via XGal

Xgal staining protocol causes blue staining to indicate the activity of β-galactosidase, the affected enzyme in GM1 gangliosidosis. In the GM1SV3 cell line, negligible blue color is present (Figure 4A). NSV3 cell line stained with Xgal presented with a blue color indicative of Bgal (Figure 4B). Both results are consistent with diseased tissue slices of GM1 feline brain and spinal cord (Figure 4C).

**Figure 4.**
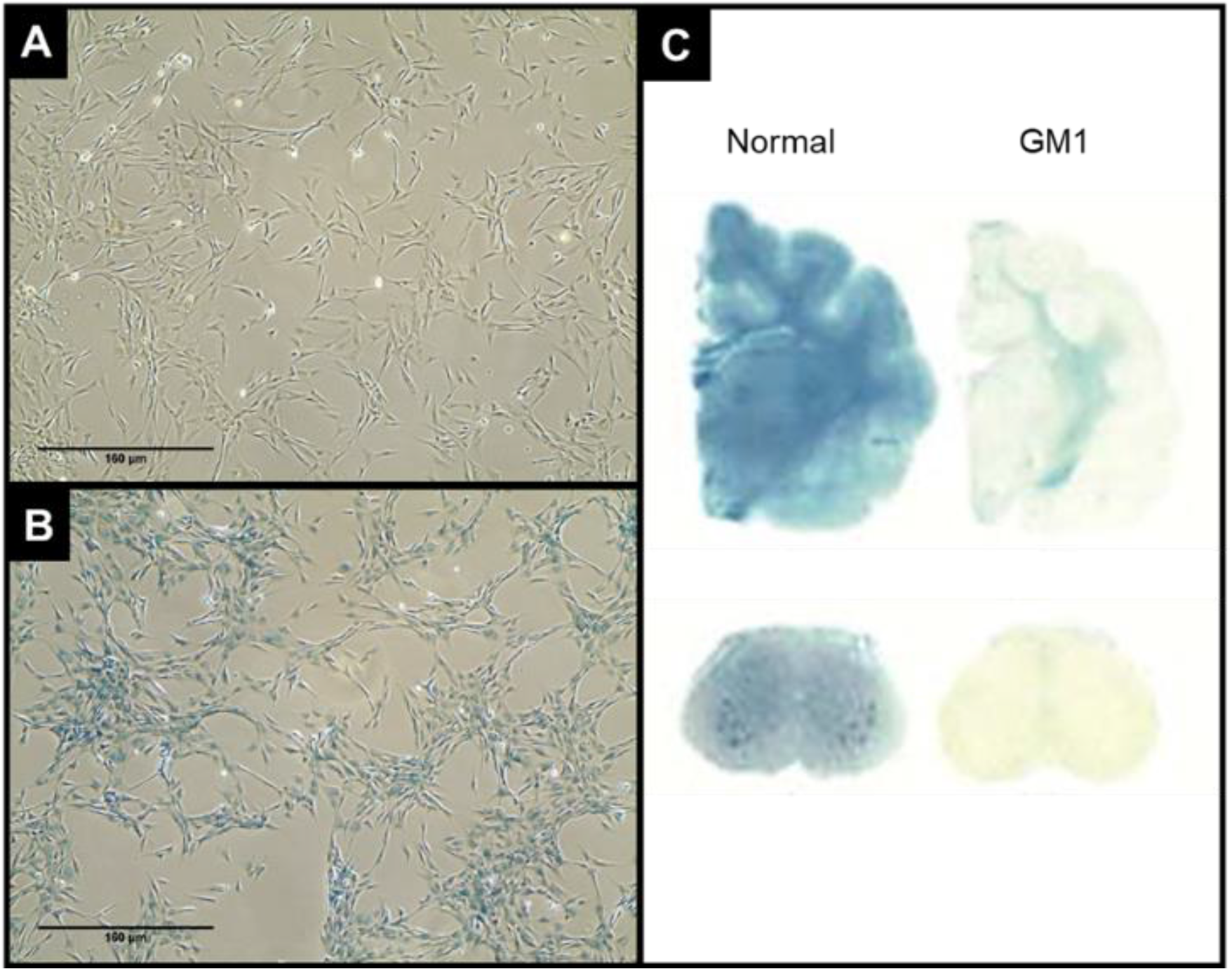
Xgal staining results of NSV3 (A) and GM1SV3 (B) cellular lines, compared to (C) published results from GM1-affected felines^49^

### 3.4. NSV3 Starvation Immunofluorescence and Quantification

Starvation is performed to induce autophagic failure^24–26^. Increasing fluorescent intensities appear to correlate with increasing starvation times of NSV3 cells. After 2 and 4 hour starvation times, NSV3 cells present with low red and green intensities of autophagosomes and lysosomes respectively, in addition to yellow areas in the overlays. After starvation times of 6 and 12 hours, fluorescence intensities of red and green independently begin to increase. Finally, after 24 and 48 hour starvation, NSV3 cells present with very intense areas of red and green with minimal to no areas of yellow (Figure 5). These observations are corroborated by image analysis measuring the percent co-localization per cell (Fig 6). Notably, after 48 hours of starvation, with a co-localization percentage of 6.5 ±7.5%, NSV3 behavior is no longer statistically different from GM1SV3 cells.

**Figure 5.**
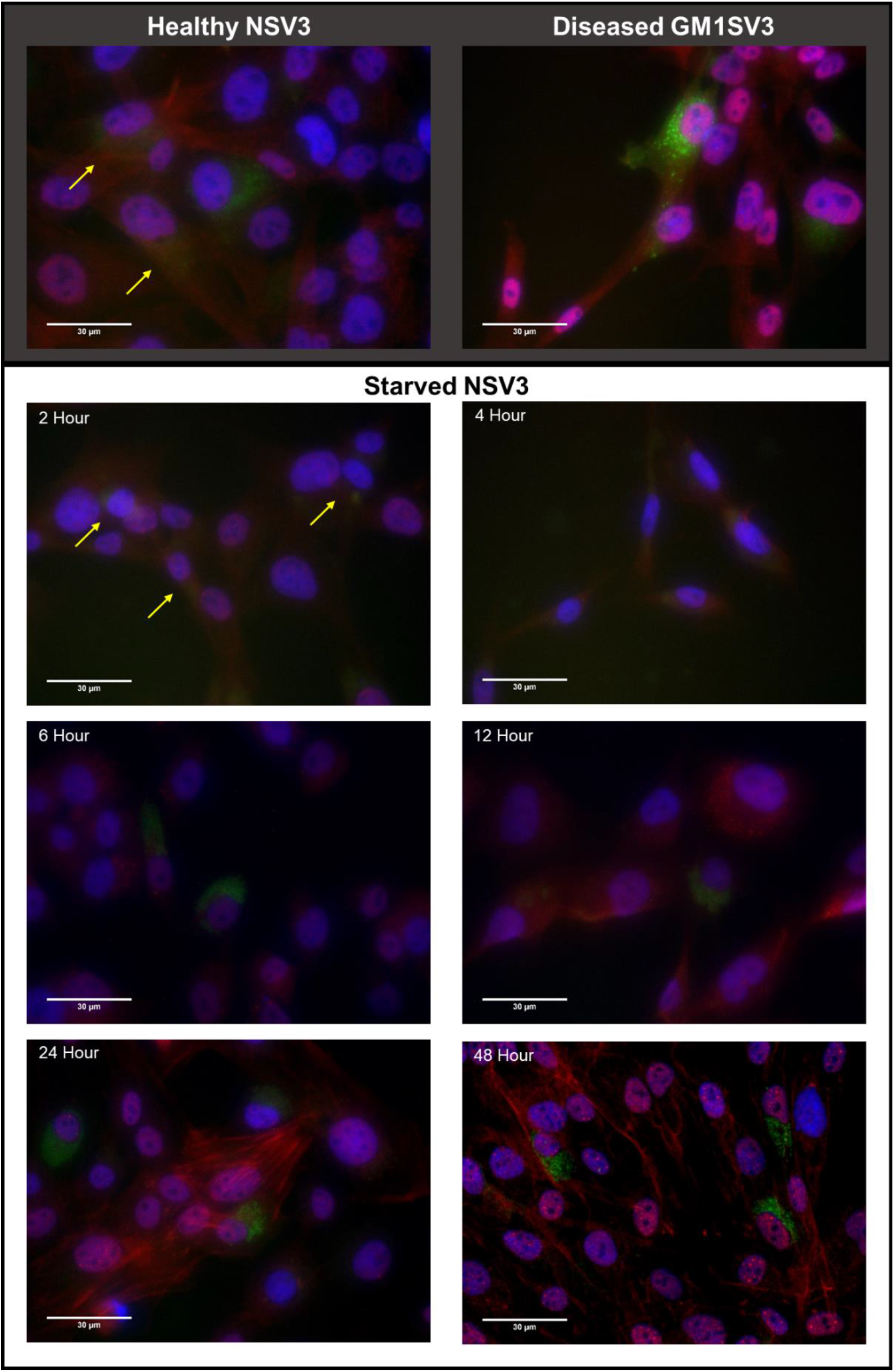
Behavior of Starved NSV3 cells at various time points. These results show that starvation causes otherwise healthy cells (NSV3) to appear more like diseased cells (GM1SV3). Left panel is provided for comparison.

**Figure 6.**
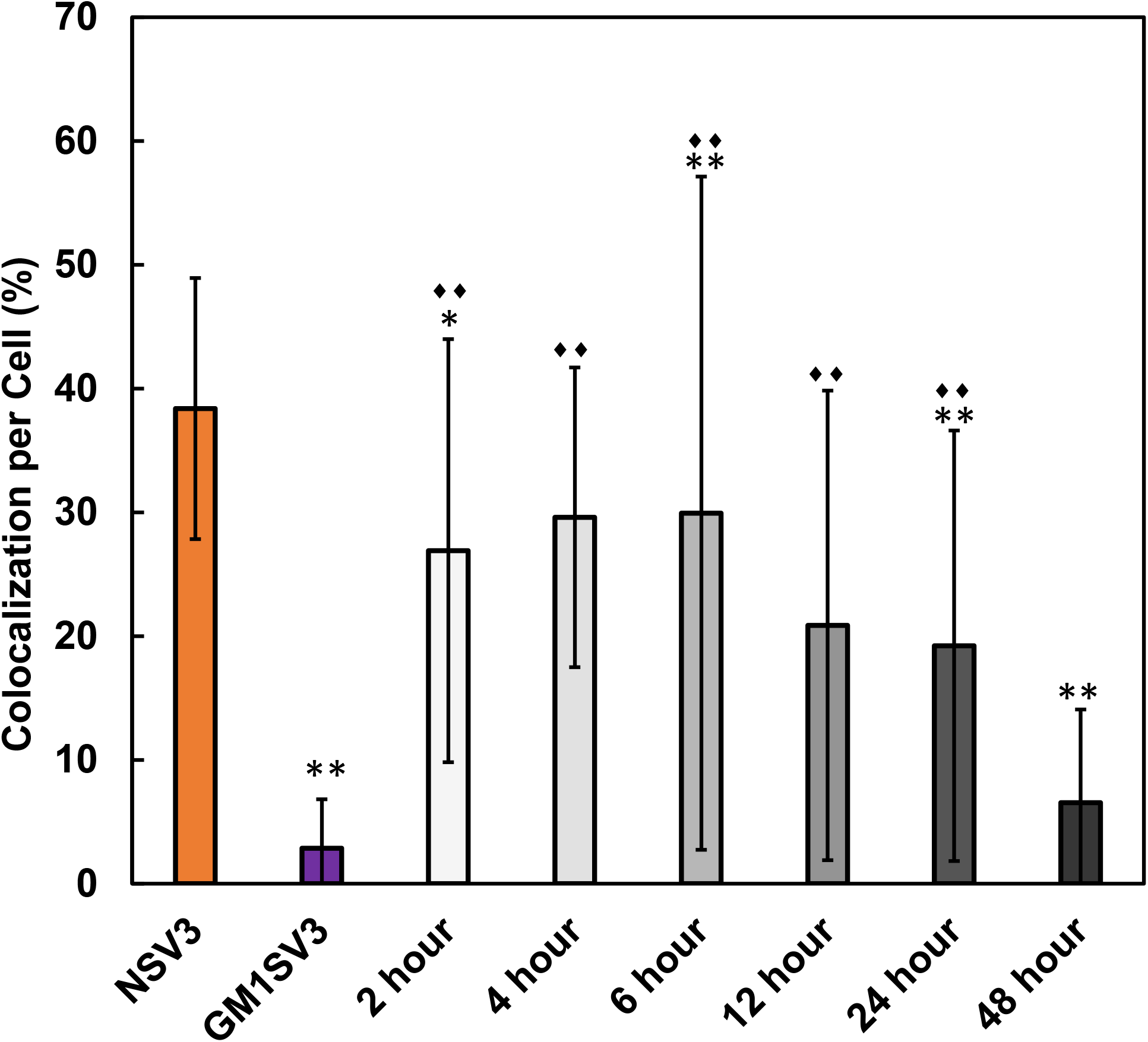
Quantification of Co-localization Observed in NSV3, GM1SV3, and starved NSV3 cells at various time points. Star (*) denotes significantly different from NSV3, with * = p<0.05, and ** = p< 0.025. Diamond (◊) denotes statistically different from GM1SV3, with ◊◊ = p < 0.025.

### 3.5. Enzyme Activities

Standard enzyme assay analysis^27,28^ indicates increased levels of lysosomal hydrolases mannosidase and hexosaminidase A in GM1SV3 cells compared to NSV3 cells. Presented in Figure 7 as fold normal enzyme activities demonstrate the deviation between GM1SV3 and NSV3 cellular lysosomal enzyme activities, with larger numbers above a fold normal of one indicating upregulation. Furthermore, enzyme upregulation trends with increased starvation time. Values corresponding to results are found in Table 1.

**Figure 7.**
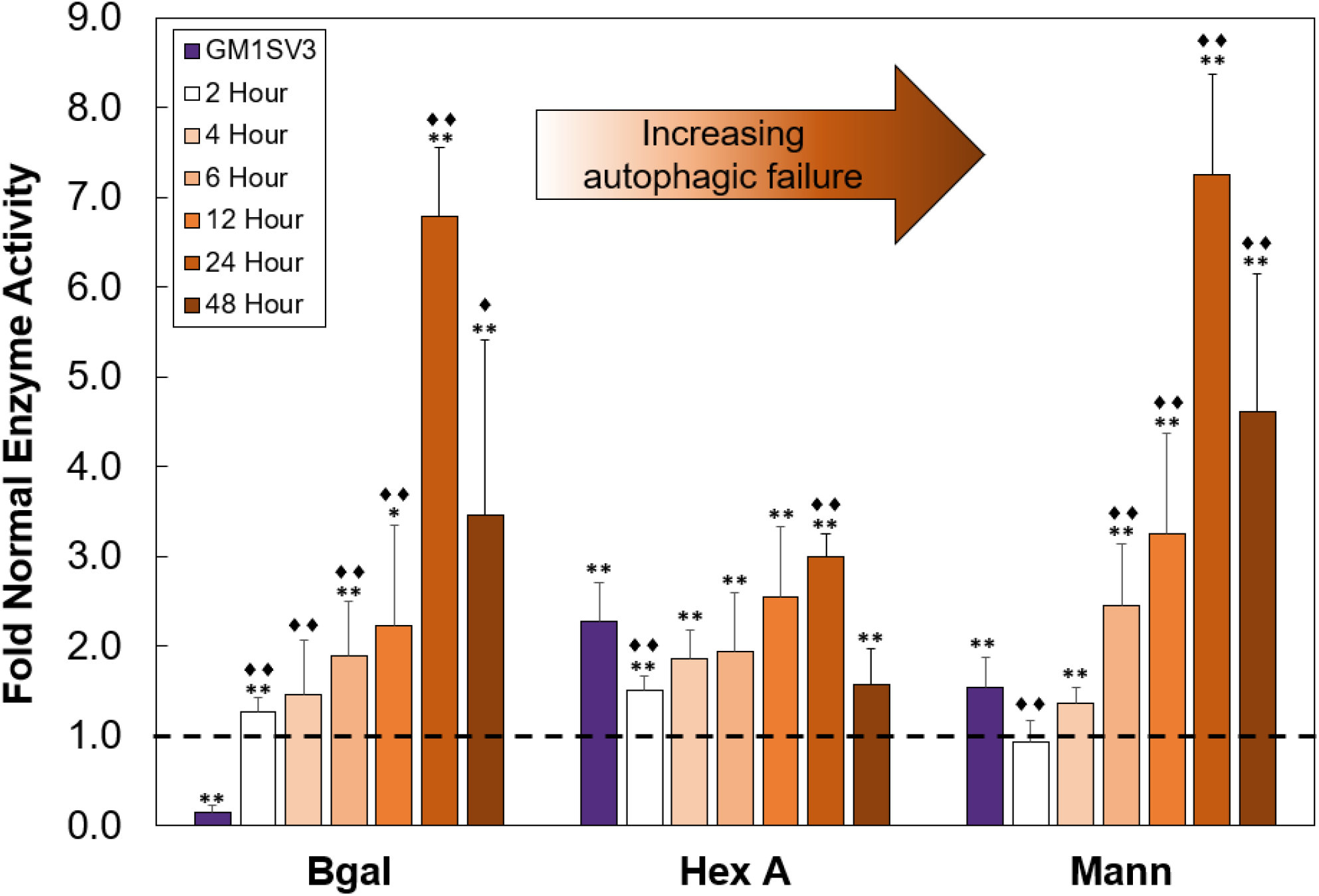
Fold normal enzyme activities of GM1SV3 and NSV3 starvation compared to healthy NSV3 of β-gal, Hex A, and Mann. Star (*) denotes significantly different from NSV3, with * = p<0.05, and ** = p< 0.025. Diamond (◊) denotes statistically different from GM1SV3, with ◊◊ = p < 0.025.

## 4. Discussion

### 4.1. Characterization of NSV3 and GM1SV3 Cell Lines

Phenotypically, NSV3 and GM1SV3 cells appear to be the same. These cell lines also match with enzyme activity levels expected based on *in vivo* results from GM1 gangliosidosis affected felines (Figures 3 and 4). Because of this, GM1SV3 cells appear to be valid as a model of GM1 gangliosidosis. It was also observed that NSV3 and GM1SV3 show differing levels of autophagic activity.

Greater intensity of red and green in GM1SV3 cells signifies an increased presence of autophagosomes and lysosomes independently. Yellow in overlay images indicates that there is co-localization of lysosomes and autophagosomes. This yellow present in NSV3 images mean that there is likely normal fusion of the organelles to form autolysosomes. The lack of presence of yellow in GM1SV3 samples indicates that there is impaired fusion and that normal autophagy is disrupted. This was expected as similar imaging results were seen in a mouse model of Multiple Sulfatase Deficiency (MSD)^23^, another LSD, with a decrease in co-localization of around 40% compared to wild-type animals. However, results presented here were more dramatic than these, with an almost 90% decrease in co-localization. This difference could be explained by the differences occurring between diseases and disease models, with MSD co-localization studies using embryonic fibroblasts of MSD-affected mice and this study using skin fibroblasts of GM1-ganglisodisosis affected felines. Also previously, an increase of LC3-II, an autophagosome specific marker, in GM1 Gangliosidosis^29,30^ has been observed. This was observed by fluorescent imaging of LC3-II in βgal −/− mice when compared to wild type. The LC3-II intensity was much greater in the βgal −/− mice samples. This is also shown in our GM1SV3 model when tagging autophagosome LC3-II with LC3 antibody. In the model of GM1SV3, the red intensity tagging autophagosomes was greater than NSV3. This observed increase of LC3-II indicates that there is a buildup of autophagosomes that are not fusing with lysosomes to form the autophagosome complex.

### 4.2. NSV3 Starvation Immunofluorescence and Quantification

Placing NSV3 cells under starvation conditions induced impaired autophagy. This is shown by the increasing red and green intensities in immunofluorescent images. This indicates that there is an increase in lysosomes and autophagosomes. The presence of yellow, or co-localization, also decreases with increased starvation showing that fusion of autophagosomes and lysosomes begins to become impaired after 6 hours (Figures 5 and 6). A similar phenomenon was observed in a model of MSD^23^, where starving mouse embryonic fibroblasts affected with MSD leading to even further autophagic failure denoted by a further decrease in co-localization of around 60% compared to wild-type mice. The MSD study was looking at the end-point effect of starvation on autophagy, where this paper attempts to study the impact of varying levels of autophagic failure on enzyme activity levels. Seeing that starvation further impairs autophagy in other lysosomal storage disorders helps to validate our hypothesis that levels of autophagic failure can be mimicked by starvation.

### 4.3. Enzyme Activities

An increase of lysosomal enzymes is seen as NSV3 cells are starved at different time points. Most notably as starvation time is increased, enzyme activity levels begin to reach and exceed activity levels of diseased GM1SV3 cells. There is a correlation between the induction of impaired autophagy and the increase of lysosomal hydrolases. It is unlikely that this increase in enzymes is due to longer potential growth times, as when starved of FBS, no growth factors are available for the cells to proliferate. To the best of our knowledge, this autophagy/enzyme relationship has not yet been observed in other models.

## 5. Conclusions

The data presented here suggests a connection between impaired autophagy and enzyme upregulation in GM1 Gangliosidosis. Research has consistently shown that enzyme upregulation occurs in GM1 gangliosidosis-affected animals in tandem with limited to no β-galactosidase production. As such, these activity levels correspond to disease. Combining this with our newly collected data indicates that there is a potential link between early stage disease, which relates to autophagosome accumulation (Figure 1), and this hydrolase regulation. Autophagic failure, although known to be an early effect of neurodegeneration, is not measurable *in situ.* The results presented here identify that hydrolase activity may be a biomarker indicative of autophagic failure, and thus neurodegeneration. Enzyme activity levels, with upregulation during disease, can be used as a diagnostic, non-invasive, biomarker to detect and track the disease in early cellular onset of neurodegeneration: the impairment of the autophagic process. Furthermore, this discovery indicates that impaired autophagy could be used as a target for treatment. To support this idea, two novel drug candidates, amodiaquine dihydrochloride dehydrate and thiethylperazine malate, that have the potential to treat GM1 Gangliosidosis target and activate autophagy^31^.

On a cellular level, neurodegeneration leads to increased autophagy in many other neurodegenerative conditions^4,7,35–40,8,15,23,25,30,32–34^. However, current methods of monitoring disease progression are limited to sacrificial, endpoint measurements. Although multiple neurodegenerative diseases^22,41–48^ are characterized by the upregulation of lysosomal hydrolases early in neurodegeneration, the relationship between enzyme upregulation and autophagy remains mostly unknown. Knowing this, the conclusions drawn and the methods developed in this paper have the potential to set a basis of information into determining a universal biomarker for the diagnosis and progression of neurodegenerative diseases.

## Acknowledgements

Cell lines provided by Dr. Douglas Martin, Auburn University.

## Funding Details

This project was funded in part by the National Institutes of Health Project number 5P20GM103499-19 through the Developmental Research Project Program. This work was supported in part by the National Science Foundation EPSCoR Program under NSF Award # OIA-1655740. Any Opinions, findings and conclusions or recommendations expressed in this material are those of the author(s) and do not necessarily reflect those of the National Science Foundation.

## Disclosure Statement

None to disclose.

---

**Supplemental Figure 1.**
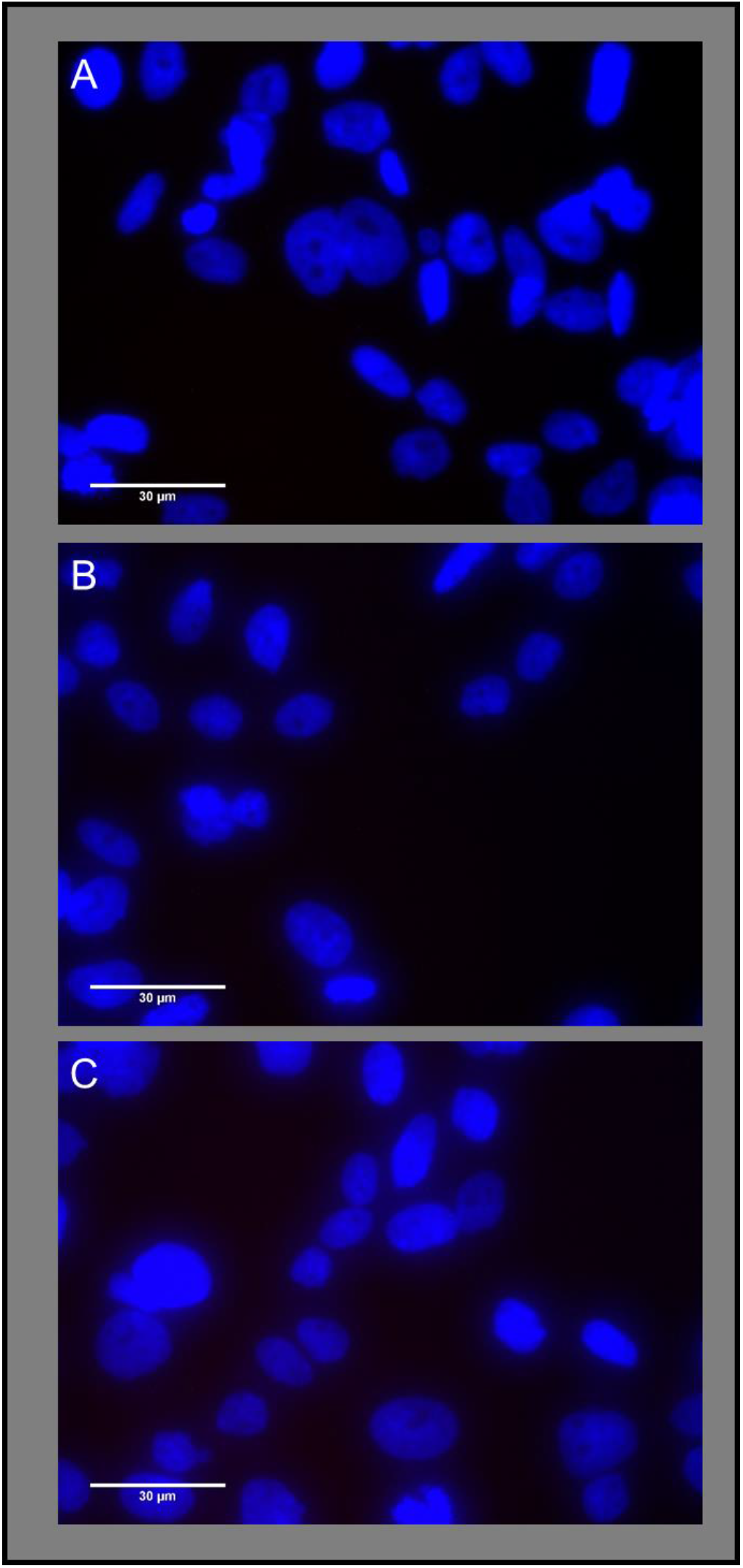
(A) Control with no antibodies present, (B) primary antibody only, (C) secondary antibody only.

## Notes

### Competing Interest Statement

The authors have declared no competing interest.

## References

1. Organization WH. Neurological disorders: public health challenges. 2006.

2. Klein AD, Mazzulli JR. Is Parkinson’s disease a lysosomal disorder? Brain 2018;

3. Niimi Y, Ito S, Mizutani Y, Murate K, Shima S, Ueda A, Satake W, Hattori N, Toda T, Mutoh T. Altered regulation of serum lysosomal acid hydrolase activities in Parkinson’s disease: A potential peripheral biomarker? Parkinsonism Relat Disord 2018;:0–1.

4. Wang C, Telpoukhovskaia M, Bahr B, Chen X, Gan L. Endo-lysosomal dysfunction: a converging mechanism in neurodegenerative diseases. Curr Opin Neurobiol 2018; 48:52–8.

5. Park JT, Lee Y-S, Cho KA, Park SC. Adjustment of the lysosomal-mitochondrial axis for control of cellular senescence. Ageing Res Rev 2018; 47:176–82.

6. Lie PPY, Nixon RA. Lysosome trafficking and signaling in health and neurodegenerative diseases. Neurobiol. Dis.2019; 122.

7. Ren H, Wang G. Autophagy and Lysosome Storage Disorders. Adv Exp Med Biol 2020; 1207:87–102.

8. Levine B, Yuan J. Autophagy in cell death: An innocent convict? J Clin Invest 2005; 115:2679–88.

9. Beck M. The Link Between Lysosomal Storage Disorders and More Common Diseases. J Inborn Errors Metab Screen 2016; 4:232640981668276.

10. Tiribuzi R, Orlacchio A, Crispoltoni L, Maiotti M, Zampolini M, De Angelis M, Mecocci P, Cecchetti R, Bernardi G, Datti A, et al. Lysosomal β-galactosidase and β-hexosaminidase activities correlate with clinical stages of dementia associated with Alzheimer’s disease and type 2 diabetes mellitus. J Alzheimer’s Dis 2011; 24:785–97.

11. Parnetti L, Chiasserini D, Persichetti E, Eusebi P, Varghese S, Qureshi MM, Dardis A, Deganuto M, De Carlo C, Castrioto A, et al. Cerebrospinal fluid lysosomal enzymes and alpha-synuclein in Parkinson’s disease. Mov Disord 2014; 29:1019–27.

12. Rocha EM, De Miranda B, Sanders LH. Alpha-synuclein: Pathology, mitochondrial dysfunction and neuroinflammation in Parkinson’s disease. Neurobiol Dis 2018; 109:249–57.

13. Ferrari R. Mitochondria and the heart. Eur Heart J 2011; 32:2911–8.

14. Oakes JA, Davies MC, Collins MO. TBK1: a new player in ALS linking autophagy and neuroinflammation. Mol Brain 2017; 10:1–10.

15. Tessitore A, Pirozzi M, Auricchio A. Abnormal autophagy, ubiquitination, inflammation and apoptosis are dependent upon lysosomal storage and are useful biomarkers of mucopolysaccharidosis VI. Pathogenetics 2009; 2:4.

16. Vanderstichele H, Bibl M, Engelborghs S, Le Bastard N, Lewczuk P, Molinuevo JL, Parnetti L, Perret-Liaudet A, Shaw LM, Teunissen C, et al. Standardization of preanalytical aspects of cerebrospinal fluid biomarker testing for Alzheimer’s disease diagnosis: A consensus paper from the Alzheimer’s Biomarkers Standardization Initiative. Alzheimer’s Dement 2012; 8:65–73.

17. El Kadmiri N, Said N, Slassi I, El Moutawakil B, Nadifi S. Biomarkers for Alzheimer Disease: Classical and Novel Candidates’ Review. Neuroscience [Internet] 2018; 370:181–90. Available from: https://doi.org/10.1016/j.neuroscience.2017.07.017

18. Gaenslen A, Unmuth B, Godau J, Liepelt I, Di Santo A, Schweitzer KJ, Gasser T, Machulla HJ, Reimold M, Marek K, et al. The specificity and sensitivity of transcranial ultrasound in the differential diagnosis of Parkinson’s disease: a prospective blinded study. Lancet Neurol 2008; 7:417–24.

19. Michell AW, Lewis SJG, Foltynie T, Barker RA. Biomarkers and Parkinson’s disease. Brain 2004; 127:1693–705.

20. Brunetti-Pierri N, Scaglia F. GM1 gangliosidosis: review of clinical, molecular, and therapeutic aspects. Mol Genet Metab [Internet] 2008 [cited 2013 Oct 10]; 94:391–6. Available from: http://www.ncbi.nlm.nih.gov/pubmed/18524657

21. Sandhoff K, Harzer K. Gangliosides and gangliosidoses: principles of molecular and metabolic pathogenesis. J Neurosci [Internet] 2013; 33:10195–208. Available from: http://www.ncbi.nlm.nih.gov/pubmed/23785136

22. McCurdy VJ, Johnson AK, Gray-Edwards HL, Randle AN, Brunson BL, Morrison NE, Salibi N, Johnson JA, Hwang M, Beyers RJ, et al. Sustained Normalization of Neurological Disease after Intracranial Gene Therapy in a Feline Model. Sci Transl Med 2014; 6:231ra48.

23. Settembre C, Fraldi A, Jahreiss L, Spampanato C, Venturi C, Medina D, de Pablo R, Tacchetti C, Rubinsztein DC, Ballabio A. A block of autophagy in lysosomal storage disorders. Hum Mol Genet 2008; 17:119–29.

24. Ganley IG, Wong PM, Gammoh N, Jiang X. Distinct Autophagosomal-Lysosomal Fusion Mechanism Revealed by Thapsigargin-Induced Autophagy Arrest. Mol Cell 2011; 42:731–43.

25. Komatsu M, Waguri S, Chiba T, Murata S, Iwata J, Tanida I, Ueno T, Koike M, Uchiyama Y, Kominami E, et al. Loss of autophagy in the central nervous system causes neurodegeneration in mice. Nature 2006; 441:880–4.

26. Mizushima N, Yamamoto A, Matsui M, Yoshimori T, Ohsumi Y. In Vivo Analysis of Autophagy in Response to Nutrient Starvation Using Transgenic Mice Expressing a Fluorescent Autophagosome Marker. Mol Biol Cell 2003; 15:1101–11.

27. Kelly JM, Gross AL, Martin DR, Byrne ME. Polyethylene glycol-b-poly(lactic acid) polymersomes as vehicles for enzyme replacement therapy. Nanomedicine 2017; 12:2591–606.

28. Martin DR, Rigat BA, Foureman P, Varadarajan GS, Hwang M, Krum BK, Smith BF, Callahan JW, Mahuran DJ, Baker HJ. Molecular consequences of the pathogenic mutation in feline GM1 gangliosidosis. Mol Genet Metab 2008; 94:212–21.

29. Lieberman AP, Puertollano R, Raben N, Slaugenhaupt S, Walkley SU, Ballabio A. Autophagy in lysosomal storage disorders. Autophagy 2012; 8:719–30.

30. Takamura A, Higaki K, Kajimaki K, Otsuka S, Ninomiya H, Matsuda J, Ohno K, Suzuki Y, Nanba E. Enhanced autophagy and mitochondrial aberrations in murine GM1-gangliosidosis. Biochem Biophys Res Commun 2008; 367:616–22.

31. Kajihara R, Numakawa T, Odaka H, Yaginuma Y, Fusaki N, Okumiya T, Furuya H, Inui S, Era T. Novel Drug Candidates Improve Ganglioside Accumulation and Neural Dysfunction in GM1 Gangliosidosis Models with Autophagy Activation. Stem Cell Reports 2020; 14:909–23.

32. Koike M, Shibata M, Waguri S, Yoshimura K, Tanida I, Kominami E, Gotow T, Peters C, von Figura K, Mizushima N, et al. Participation of autophagy in storage of lysosomes in neurons from mouse models of neuronal ceroid-lipofuscinoses (Batten disease). Am J Pathol 2005; 167:1713–28.

33. Raben N, Schreiner C, Baum R, Takikita S, Xu S, Xie T, Myerowitz R, Komatsu M, Van der Meulen JH, Nagaraju K, et al. Suppression of autophagy permits successful enzyme replacement therapy in a lysosomal storage disorder--murine Pompe disease. Autophagy 2010; 6:1078–89.

34. Green DR, Galluzzi L, Kroemer G. Mitochondria and the Autophagy-Inflammation-Cell Death Axis in Organismal Aging. Science (80-) 2011; 333:1109–12.

35. Guo F, Liu X, Cai H, Le W. Autophagy in neurodegenerative diseases: Pathogenesis and therapy. Brain Pathol 2017;

36. Kaizuka T, Morishita H, Hama Y, Tsukamoto S, Matsui T, Toyota Y, Kodama A, Ishihara T, Mizushima T, Mizushima N. An Autophagic Flux Probe that Releases an Internal Control. Mol Cell 2016; 64:835–49.

37. Ballabio A, Gieselmann V. Lysosomal disorders: from storage to cellular damage. Biochim Biophys Acta 2009; 1793:684–96.

38. Vitner EB, Platt FM, Futerman AH. Common and uncommon pathogenic cascades in lysosomal storage diseases. J Biol Chem 2010; 285:20423–7.

39. Martins C, Hulkova H, Dridi L, Dormoy-Raclet V, Grigoryeva L, Choi Y, Langford-Smith A, Wilkinson FL, Ohmi K, DiCristo G, et al. Neuroinflammation, mitochondrial defects and neurodegeneration in mucopolysaccharidosis III type C mouse model. Brain 2015; 138:336–55.

40. Hara T, Nakamura K, Matsui M, Yamamoto A, Nakahara Y, Suzuki-Migishima R, Yokoyama M, Mishima K, Saito I, Okano H, et al. Suppression of basal autophagy in neural cells causes neurodegenerative disease in mice. Nature 2006; 441:885–9.

41. Martinez-Vicente M, Cuervo AM. Autophagy and neurodegeneration: when the cleaning crew goes on strike. Lancet Neurol 2007; 6:352–61.

42. Bendiske J, Bahr B a. Lysosomal activation is a compensatory response against protein accumulation and associated synaptopathogenesis--an approach for slowing Alzheimer disease? J Neuropathol Exp Neurol 2003; 62:451–63.

43. Orr ME, Oddo S. Autophagic/lysosomal dysfunction in Alzheimer’s disease. Alzheimers Res Ther 2013; 5:53.

44. Cataldo a M, Hamilton DJ, Barnett JL, Paskevich P a, Nixon R a. Properties of the endosomal-lysosomal system in the human central nervous system: disturbances mark most neurons in populations at risk to degenerate in Alzheimer’s disease. J Neurosci 1996; 16:186–99.

45. Pentchev PG, Gal AE, Booth AD, Omodeo-Sale F, Fouks J, Neumeyer BA, Quirk JM, Dawson G, Brady RO. A lysosomal storage disorder in mice characterized by a dual deficiency of sphingomyelinase and glucocerebrosidase. Biochim Biophys Acta 1980; 619:669–79.

46. Lloyd-Evans E, Morgan AJ, He X, Smith DA, Elliot-Smith E, Sillence DJ, Churchill GC, Schuchman EH, Galione A, Platt FM. Niemann-Pick disease type C1 is a sphingosine storage disease that causes deregulation of lysosomal calcium. Nat Med 2008; 14:1247–55.

47. Kopacek J, Sakaguchi S, Shigematsu K, Nishida N, Atarashi R, Nakaoke R, Moriuchi R, Niwa M, Katamine S. Upregulation of the genes encoding lysosomal hydrolases, a perforin-like protein, and peroxidases in the brains of mice affected with an experimental prion disease. J Virol 2000; 74:411–7.

48. Butler D, Nixon RA, Bahr BA. Potential compensatory responses through autophagic/lysosomal pathways in neurodegenerative diseases. Autophagy 2006; 2:234–7.

49. Martin DR, Cox NR, Morrison NE, Kennamer DM, Peck SL, Dodson AN, Gentry AS, Griffin B, Rolsma MD, Baker HJ. Mutation of the GM2 activator protein in a feline model of GM2 gangliosidosis. Acta Neuropathol 2005; 110:443–50.

50. McCurdy VJ, Rockwell HE, Arthur JR, Bradbury AM, Johnson AK, Randle AN, Brunson BL, Hwang M, Gray-Edwards HL, Morrison NE, et al. Widespread correction of central nervous system disease after intracranial gene therapy in a feline model of Sandhoff disease. Gene Ther 2015; 22:181–9.

